# Voltage imaging reveals the emergence of population activity in the spinal cord

**DOI:** 10.1101/2023.05.25.540669

**Authors:** Asuka Shiraishi, Ayane Hayashi, Narumi Fukuda, Mari Hishinuma, Hiroaki Miyazawa, Sachiko Tsuda

**Affiliations:** Division of Life Science, Graduate School of Science and Engineering, Saitama University, 255 Shimo-Okubo, Sakura-ku, Saitama City, Saitama 338-8570, Japan; Department of Regulatory Biology, Faculty of Science, Saitama University, 255 Shimo-Okubo, Sakura-ku, Saitama City, Saitama 338-8570, Japan

## Abstract

One of the central questions in neural development is how individual neurons assemble functional networks. To address this, it is essential to elucidate how coordinated activity emerges during development. However, tracking the functional maturation of neuronal populations over time remains challenging, as it requires long-term, non-invasive monitoring of membrane potential dynamics. Here, we developed a voltage imaging approach for zebrafish embryos using genetically encoded voltage indicators (GEVIs), enabling fast, direct and cell-type-specific measurements of membrane potentials from defined neuronal populations in a non-invasive manner. Using this approach, we detected coordinated voltage changes in spinal motor neurons with high spatiotemporal resolution. Depolarization and hyperpolarization events were observed at the population, single-cell, and subcellular levels. Notably, long-term voltage imaging revealed the early emergence and progressive maturation of membrane potential dynamics, characterized by increased firing rate, coupling strength and axonal outgrowth. This optical approach constitutes a significant advancement in the study of neural development, providing a powerful tool for investigating the spatiotemporal dynamics of neuronal populations in vivo.

## Introduction

The nervous system is composed of a vast number of neurons that perform diverse functions. How are these neurons generated and how do they communicate with each other to form functional networks? To better understand the function and development of the nervous system, it is crucial to examine the dynamic behavior of neuronal populations and how these dynamics evolve over time. However, such investigations are often hampered by the huge number of neurons involved and the invasiveness of conventional electrophysiological recording techniques.

Optical methods provide a powerful alternative, as light facilitates non-invasive access to large populations of neurons, which is difficult with electrophysiological recordings that insert electrodes into target tissues as single-point detectors. Voltage imaging enables fast, direct, and simultaneous detection of membrane potentials across multiple cells, providing an excellent system for understanding the dynamic behavior of neuronal populations [1–3]. Its high speed and directness allow the detection of action potentials, hyperpolarization, and subthreshold responses, which are difficult to examine using calcium imaging, partly because of the slow kinetics of the indicators and the dynamics of calcium [4, 5]. Various voltage-sensitive dyes have been developed to improve brightness, signal-to-noise ratio, and spectral range, thereby advancing our understanding of network dynamics in the brain [6–10]. More recently, genetically encoded voltage indicators (GEVIs) have been developed, allowing for cell-type-specific voltage imaging [11–13]. These GEVIs include the ones based on voltage-sensitive phosphatase domains [14–17], as well as microbial rhodopsin-based sensors [18–21]. Although GEVIs have been successfully applied to investigate brain and cardiac function [22–29], their application to developmental studies of the nervous system remains limited. This is largely due to the technical challenges of achieving long-term, non-invasive voltage imaging in developing tissues.

One of the earliest neural structures to develop is the spinal cord, which serve as the core of rhythmic motor behaviors through central pattern generators [30]. Neurons in the spinal cord, including motor neurons, exhibit prominent population activity even at early in development. This activity drives fetal movement and is suggested to play a critical role for circuit formation [31, 32]. While electrophysiological studies have characterized the firing properties of these neurons, the spatiotemporal organization and developmental emergence of their coordinated voltage dynamics, the core of the remain to be understood [33–35].

In this study, to examine the spatiotemporal dynamics of neuronal populations and their emergence during development, we applied GEVI ArcLight to zebrafish embryos, taking advantage of their optical transparency, genetic tools, and rapid development [36, 37]. Although recent studies have employed GEVIs in zebrafish larvae primarily to explore circuit function [20, 21, 24, 25, 28], long-term developmental imaging has not yet been achieved. Here, we establish a long-term voltage imaging approach that enables the observation of voltage dynamics in neuronal populations over the course of development. This approach allowed us to reveal how coordinated voltage activity emerge during early neural development.

## Materials and Methods

### Fish lines

Wild-type zebrafish (Danio rerio) with RW genetic background or nacre line were used [38]. *Tg(elavl3:GAL4-VP16)*, *Tg(UAS:lynRFP)*, and *Tg(elavl3:GCaMP6f)* fish lines have been described previously and obtained from the National BioResource Project [24, 39]. The Tg lines were crossed with the nacre line. All procedures were performed in accordance with a protocol approved by the Saitama University Committee on Animal Research.

### DNA Construction and establishment of ArcLight transgenic line

To generate pTol2-UAS-ArcLight plasmid, ArcLight-A242 sequence was excited from the pCS2-ArcLight-A242 plasmid (Addgene) and inserted into pTol2-UAS-MCSF-polyA, in which UAS and polyA sequences were integrated into the pTol2-MCS plasmid [40]. To establish pTol2-UAS-ArcLight line, Tol2 plasmid DNA and *transposase* mRNA were co-injected into one-cell-stage embryos. Genotyping was performed using the primers for ArcLight-A242 (5′-TCGACGGTATCGATAAGCTTGC-3′,5′-CACCTCCCCCTGAACCTGAAACATA-3′).

### Transient expression of ArcLight in the spinal cord and cerebellum

For the spinal cord, 25 ng/μl pTol2-UAS-ArcLight-A242-polyA plasmid DNA was co-injected with 25 ng/μl *Tol2 transposase* mRNA into one-cell-stage embryos of *Tg(elavl3:GAL4-VP16)* embryos, which express Gal4 in neurons [39]. After the injection, the embryos were incubated at 23°C and then transferred to 28°C by 18 hour post fertilization (hpf). ArcLight-positive embryos were screened at the 18.5 to 19 hpf, followed by voltage imaging.

### Confocal imaging

Zebrafish embryos were paralyzed in 0.02% tricaine and mounted on 2% low melting-point agarose (Sigma-Aldrich). Optical sectioning was performed using an Olympus FV1000 or a Nikon A1R confocal microscope. ImageJ, NIS-Elements (Nikon), and Imaris (Bitplane) programs were used to perform the image analysis.

### Voltage imaging

Embryos from 16 to 23 hpf were paralyzed with tubocurarine (0.5 mM, Sigma-Aldrich), with their tail tips cut for better penetration of tubocurarine [41, 42]. Then, embryos were mounted on 1.8% low melting-point agarose containing 0.1-0.2 mM tubocurarine. After embedding the embryos onto 1% agarose, an extracellular solution (134 mM NaCl, 2.9 mM KCl, 2.1 mM CaCl_2_, 1.2 mM MgCl_2_, 10 mM HEPES, and 10 mM glucose, adjusted to pH 7.8 with NaOH) containing 150 mM tubocurarine was added. The agarose around the tail was removed using a surgical blade to allow efficient transfer of the drug solution. For high-speed voltage imaging, a fluorescence microscope (FN-1, Nikon) equipped with a CMOS camera (ORCA-Flash4.0, Hamamatsu Photonics) was used with a 40×/0.8 NA water-immersion lens. For most recordings, a 0.7× relay lens (NIKON) was also used. Fluorescence images were acquired using NIS-Elements (Nikon) software at approximately 16-200 Hz. High-speed confocal scanning was performed using a confocal microscope (Nikon A1R) with a 25×/1.0 NA water-immersion lens. For drug treatment, tetrodotoxin (1 μM, TOCRIS) was added to the extracellular solution using a peristaltic pump (World Precision Instruments). A thermo plate was used to keep the recording chamber at 28°C (Tokai hit).

### Image analysis

Images were analyzed using the NIS-Elements (Nikon), Fiji, and OriginPro (Lightstone) programs as described previously [41]. To extract changes in fluorescence intensity, regions of interest (ROIs) were manually set using NIS-element software (Nikon). Normalized changes in fluorescence intensity in the target regions (ROIs) were quantitatively analyzed by calculating dF/F in the ROIs (dF/F_0_=[F_(t)_–F_0_]/F_0_, where F_(t)_ and F_0_ are the fluorescence intensities at a certain time point and initial time points, respectively). If the larvae/embryos moved during the voltage imaging, the images were aligned using a Fiji program (TurboReg plugin). Denoising and bleach corrections were also applied to some data using OriginPro (Lightstone). Cross-correlation and hierarchical clustering analyses were performed using OriginPro (Lightstone). The coupling rate was calculated as the ratio of the frequency of target cell activity to that of mature cells. Among depolarizing signals with small voltage changes, those characterized by a longer time to peak and a shorter decay time were defined as "immature firing" and were distinguished from other small depolarizations.

### Electrophysiological recordings

Patch clamp recordings were conducted as described previously, using borosilicate glass pipettes at 28 ℃ (World Precision Instrument, IB150F-4WPI) [24]. Electrical responses were acquired via a Multiclamp 700B (Molecular devices) and DAQ interface (National Instruments, USB-6212 BNC) with WinWCP software (Univ. of Strathclyde), which also recorded the timing of image acquisition.

## Results

### Establishment of voltage imaging using ArcLight zebrafish embryos

To investigate voltage dynamics of neuronal populations in neuronal populations during development, we established a transgenic zebrafish line that expresses ArcLight specifically in neurons. We first established the transgenic line Tg(*UAS:ArcLight*) using the GAL4-UAS and Tol2 transposon systems [37] (Fig. 1a). To confirm ArcLight localization to cellular membranes in Tg(*UAS:ArcLight*) fish, we examined Tg(*elavl3:GAL4-VP16;UAS:ArcLight;UAS:lyn-RFP*) embryos, in which GAL4 is expressed in neurons and cellular membranes are labeled with Lyn-RFP.

**Figure 1.**
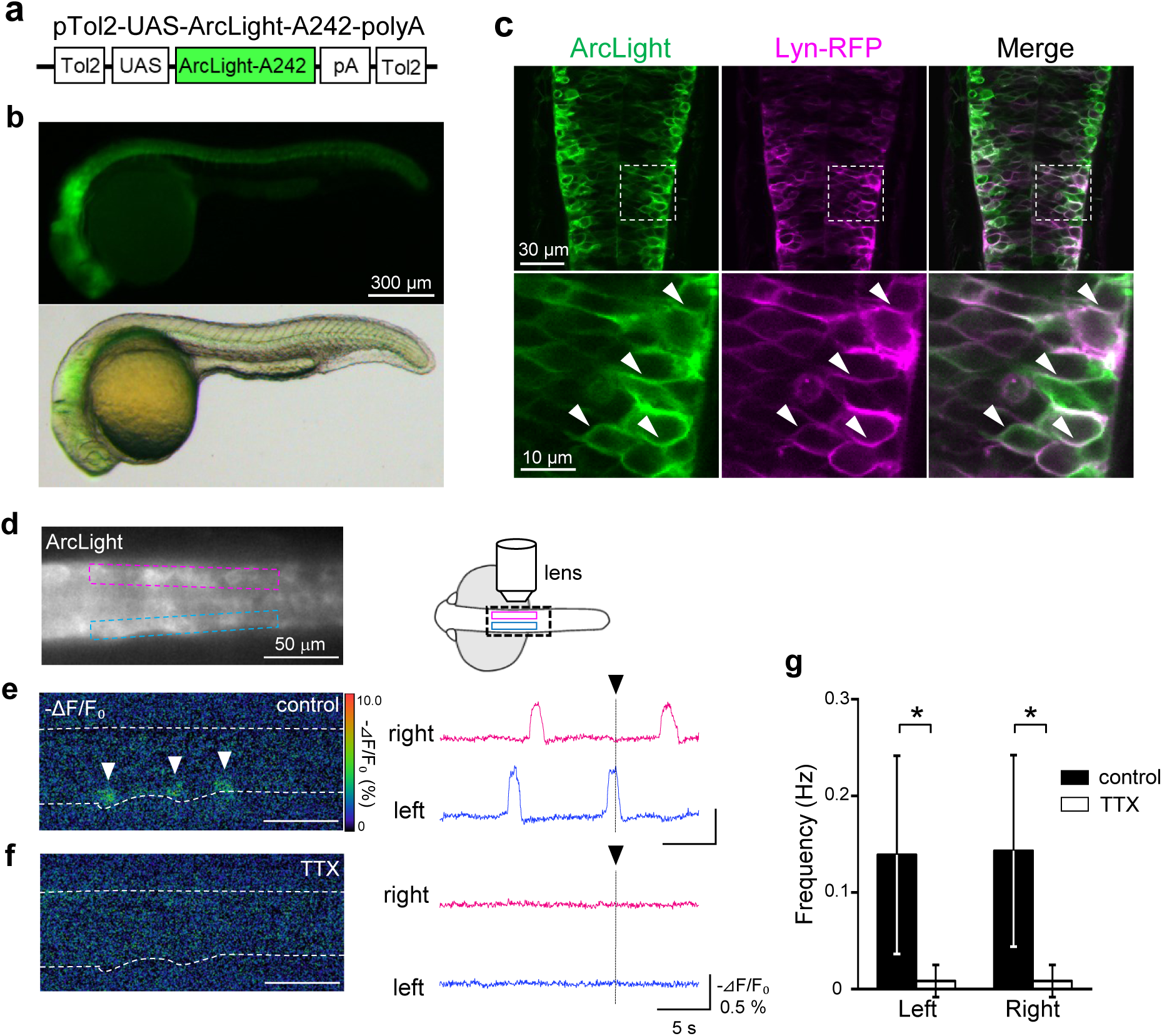
Transgenic zebrafish showing membrane-localized ArcLight and spontaneous activity in the neural tube. (a) Schematic diagram of plasmid construct for ArcLight. (b) Lateral views of ArcLight-expressing embryos at 1 dpf. ArcLight is distributed widely in the neural tube of Tg*(elavl3:GAL4-VP16;UAS:ArcLight)* fish. (c) Dorsal view of the neural tube of Tg*(elavl3:GAL4-VP16;UAS:ArcLight;UAS:lyn-RFP)* embryos at 1 dpf. ArcLight (green) is co-localized with Lyn-RFP (red: cell membranes) in the neural tube (arrowheads). Higher magnification images are shown in the lower panel. (d-f) Spontaneous activity of spinal cord neurons is detected by ArcLight imaging. (d) Dorsal view of the ventral spinal cord of Tg *(elavl3:GAL4-VP16;UAS:ArcLight-A242)* fish at 20 hpf. The rostral side is to the left, and the area between 3–8 somites is shown. A schematic diagram of the imaging system is shown on the right. Changes in the fluorescence of ArcLight (−ΔF/F_0_) are indicated by the pseudocolor scale shown at right. White arrowheads indicate the activated neurons. Regions of interest (ROIs) located between 5–7 somites are indicated by red (right side) and blue (left side) rectangular in (d). Fluorescence changes of ArcLight in the ROIs are shown on the right. The time point of the image is indicated by a black arrowhead. (f-g) The spontaneous activity in the spinal cord mostly disappeared by tetrodotoxin (TTX) treatment (control 0.14±0.10 Hz, TTX 0.01±0.02 Hz, p<0.05, right side: control 0.14±0.10 Hz, TTX 0.01±0.02 Hz, *: p<0.05, Wilcoxon signed-rank test, 6 fish). The imaged plane and ROIs are in the same positions as in (d).

At 1 dpf, ArcLight was widely observed in the neural tube and co-localized with Lyn-RFP, indicating proper membrane localization of ArcLight in neurons (Fig. 1b, c).

Spinal cord neurons exhibit spontaneous firing early in development, with coordinated activity of the neuronal populations beginning around 17 hpf [34, 43]. Consistent with this, Tg(*elavl3:GAL4-VP16;UAS:ArcLight*) embryos displayed oscillatory changes in ArcLight fluorescence in the ventral spinal cord (Fig. 1d–g, 0.14±0.10 Hz, 6 fish). Fluorescence fluctuations alternated between the left and right sides of the spinal cord, consistent with previous reports of periodic depolarization (PD) in spinal neurons [43]. This activity was largely abolished following treatment with the sodium channel blocker tetrodotoxin (TTX) (Fig. 1e-g). These results demonstrate that ArcLight can reliably detect spontaneous activity in developing spinal neurons.

We next examined whether ArcLight can resolve spinal voltage dynamics at both single-cell and population levels. As shown in Figure 2, oscillatory activity was clearly detected in individual neurons, with distinct amplitudes and temporal patterns among neighboring neurons (Fig. 2a-c). Depolarizations typically propagated sequentially along the rostrocaudal axis. In addition to large depolarizations that closely resemble PD, small membrane potential fluctuations were also observed between the PDs (Fig. 2d arrowheads). Based on their size and timing, these may be synaptic bursts or subthreshold events, which are generally only captured by electrophysiological recordings [43]. Cross-correlation analysis revealed strong positive correlations among ipsilateral neurons and negative correlations between contralateral pairs, indicating high synchrony within each side of the spinal cord (Fig. 2e). Hierarchical clustering analysis also supported this synchrony with partially disordered activity along rostrocaudal axis (Fig. 2f). The depolarization was confirmed by electrophysiological recordings (Supplementary Fig S1). To evaluate the strength of ArcLight voltage imaging, we performed spinal calcium imaging using Tg(*elavl3:GCaMP6f*) and compared its temporal resolution with ArcLight. Calcium imaging exhibited substantially lower temporal resolution and reduced variability in activity patterns (Supplementary Fig S1), highlighting the superior precision of the GEVI ArcLight in capturing neural population dynamics.

**Figure 2.**
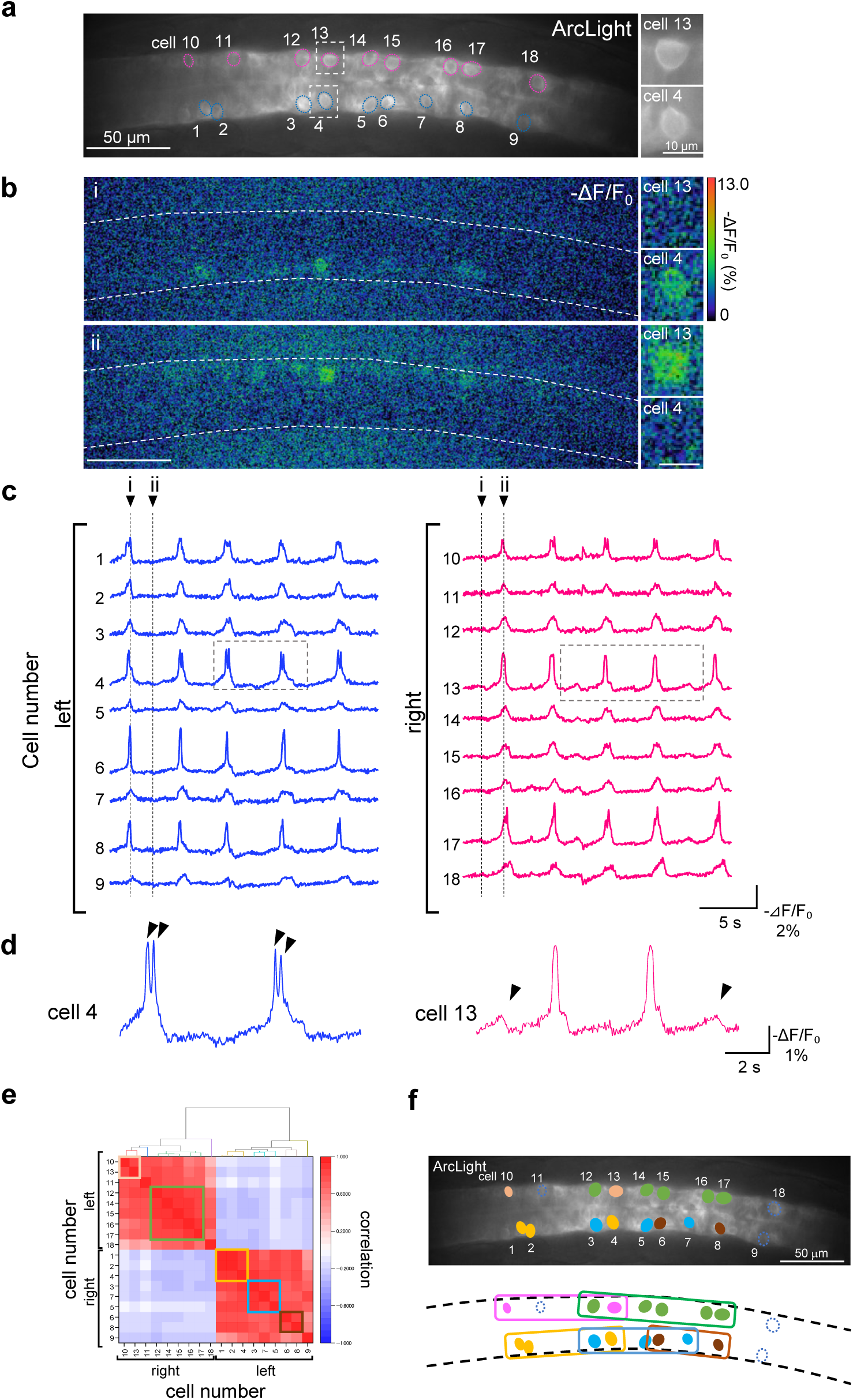
Monitoring spontaneous activity of individual cells in the spinal cord by ArcLight imaging. (a) A fluorescence image of the ventral spinal cord of Tg*(elavl3:GAL4-VP16;UAS:ArcLight)* fish at 21 hpf. ROIs are located at the 18 cells (red: right side, blue: left side). Higher magnification views are shown on the right. (b) Changes in ArcLight signal at the two time points indicated in (c). (c) Activity patterns of the 18 cells are shown. Higher magnification views of the ArcLight signal of cells 4 and 13 are shown at the bottom. (d). (e, f) Hierarchal clustering analysis of the voltage dynamics in the 18 cells shows the synchronous activity in the ipsilateral neuron pairs (e), and also several groups with higher synchrony within them (f).

### Subcellular voltage imaging of neural populations

Measuring membrane potential dynamics in subcellular compartments such as axons and dendrites is crucial for understanding neuronal information processing [44, 45]. As shown in Figure 3, ArcLight voltage imaging successfully captured depolarizations in somata as well as in proximal and distal axonal regions. Although somata and axons exhibited similar depolarization patterns, the amplitude of voltage changes was slightly smaller in axons than in somata (Fig. 3b). Axonal depolarizations were also confirmed by confocal microscopy (Supplementary Fig. S2). Furthermore, high-resolution recordings revealed voltage fluctuations and morphological dynamics in fine structures such as filopodia (Supplementary Fig. S3).

**Figure 3.**
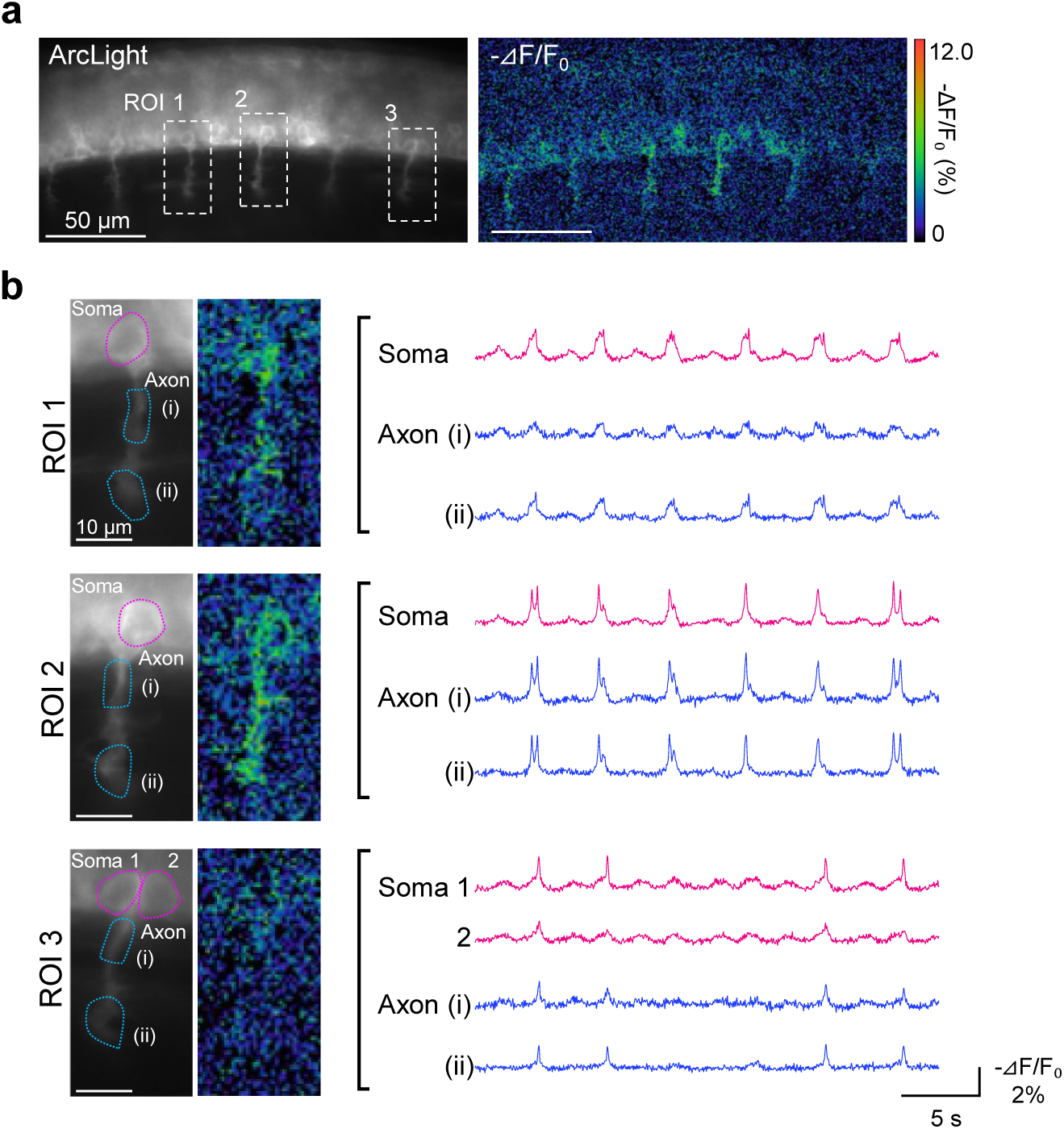
Voltage imaging of spinal cord neurons at a high spatiotemporal resolution at the cellular compartment level by ArcLight. (a) Lateral views of the spinal cord of Tg*(elavl3:GAL4-VP16;UAS:ArcLight)* fish at 20 hpf. A schematic diagram of the imaging system is shown on the right. (b) Changes in ArcLight signal are observed at soma and axons in three different ROIs shown in (a). Higher resolution observations are shown in Supplementary Fig. S2 and S3.

### Single-cell voltage imaging reveals simultaneous voltage and morphological dynamics

Spinal circuits comprise diverse neuronal types with distinct distributions, morphologies, and physiological properties [46]. To understand how these circuits develop, it is important to simultaneously monitor voltage dynamics and morphological changes. To achieve this, we used mosaic expression of ArcLight. By injecting *UAS-ArcLight* pDNA into Tg(*elavl3:GAL4-VP16*) fish, we sparsely labeled neurons, enabling high-resolution tracking of membrane potential changes and cellular morphology (Fig. 4a–c, 20 hpf, PMNs). We also captured voltage dynamics from interneurons that are otherwise difficult to identify (Fig. 4d, e). In addition to PD and synaptic burst (SB)-like activity, some neurons showed hyperpolarizing responses (Fig. 4e). Over the course of one hour, we observed notable morphological changes in these neurons, suggesting active developmental remodeling (Supplementary Fig. S4).

**Figure 4.**
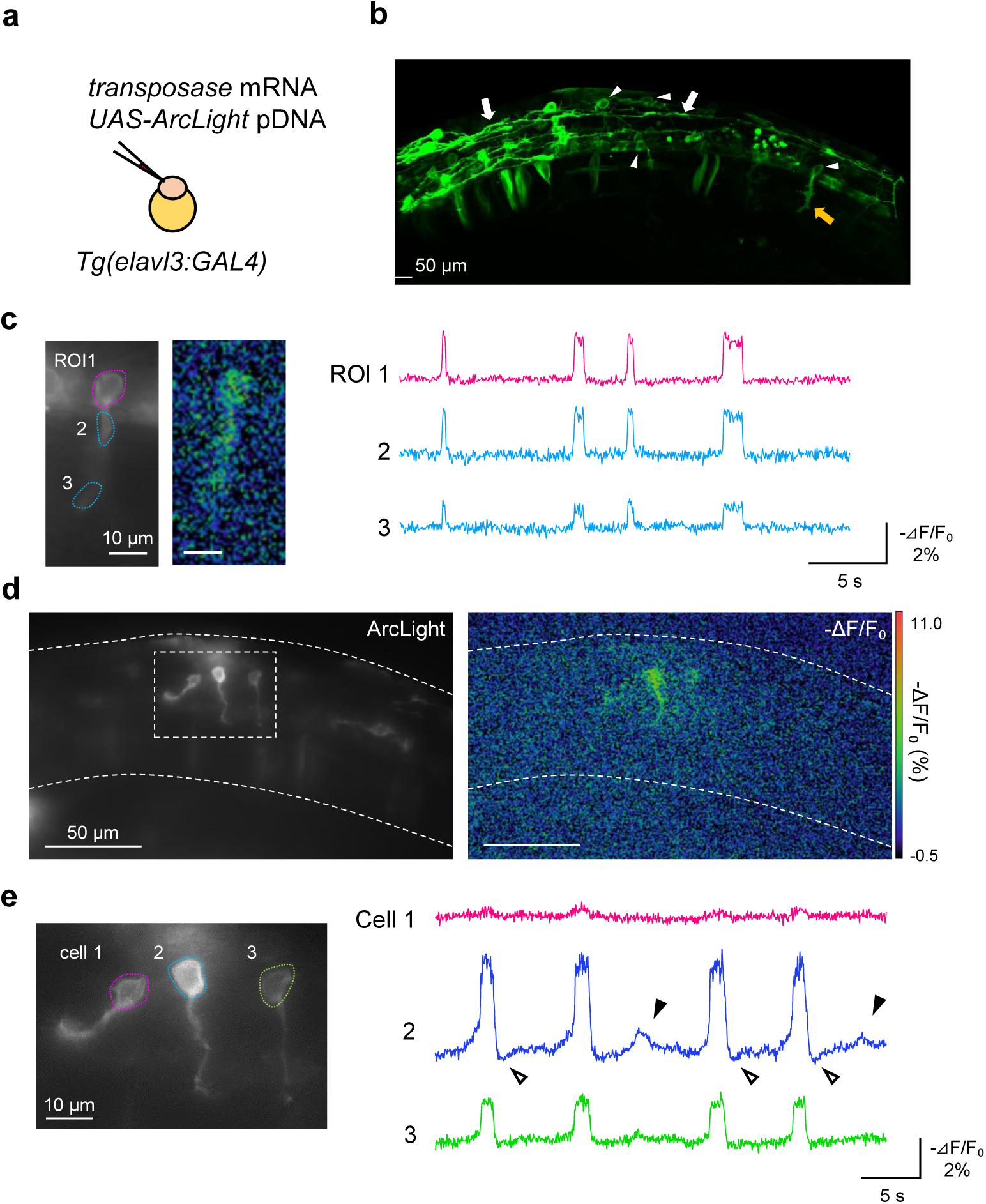
Single cell labeling of neurons by ArcLight enables simultaneous recording of voltage and morphology. (a) Schematic diagram of the co-injection experiment. (b) A stacked confocal image of the spinal cord region of the injected embryo at 20 hpf. The left side view is shown. Arrowheads and arrows indicate the soma and axons of ArcLight positive cells, respectively. (c) Voltage dynamics of a primary motor neuron are indicated by a yellow arrowhead in (b). ROIs at soma and axons are shown in red and blue, respectively. (d, e) Voltage imaging of the spinal cord interneurons. A higher magnification view is shown in (e). ArcLight signals from three cells are shown on the right. In addition to periodic depolarization, hyperpolarization (white arrowheads) and subthreshold-like signals (black arrowheads) are observed.

### Developmental changes in voltage dynamics of neuronal populations

Finally, we investigated the emergence and maturation of voltage dynamics during development. Periodic firing in the spinal cord drives coiling behavior, analogous to fetal movement in humans, which are thought to contribute to spinal circuit formation [32]. This characteristic activity appears around 17 hpf and rapidly increases in frequency and amplitude until 20 hpf, followed by a gradual decline [34]. However, how population-level coordinated activity arises during this process remains to be elucidated.

To address this, we performed long-term voltage imaging of the spinal cord along the course of development (Fig. 5). At 18 hpf, primary motor neurons (PMNs) possesses an axonal protrusion but seldom exhibited firing, instead showing small voltage fluctuations (Fig. 5c, arrowheads, cell 1, located at somite 8). These fluctuations were observed in both soma and axons (Supplementary Fig. S5) and displayed slower kinetics than mature firing (Supplementary Fig. S6). Over the next 30 minutes, PMNs gradually extended axons and increased firing frequency. By 19.5 hpf, the firing frequency reached ∼0.3 Hz (0.29±0.05 Hz). In addition to cell 1, PMN cell 2, located more caudally, showed slower axonal extension and began firing after 18.5 hpf, ultimately synchronizing with other neurons by 19.5 hpf (Fig. 5). Similar temporal maturation patterns were also observed in other PMNs with some variability (Supplementary Fig. S7). Quantitative analyses revealed that firing frequency and coordination increased in tandem with axonal extension (Fig. 6a, b). In deede, the emergence of small depolarization and matured firing strongly correlated with morphological maturation (Fig. 6c, d). Moreover, coordination among PMNs also increased alongside the maturation of their individual voltage dynamics (Fig. 6e, f). These data suggest a sequence in the emergence of coordinated activity in spinal neurons: (1) absence of depolarization, (2) small depolarizations, (3) irregularly coordinated activity, and (4) fully synchronized firing.

**Figure 5.**
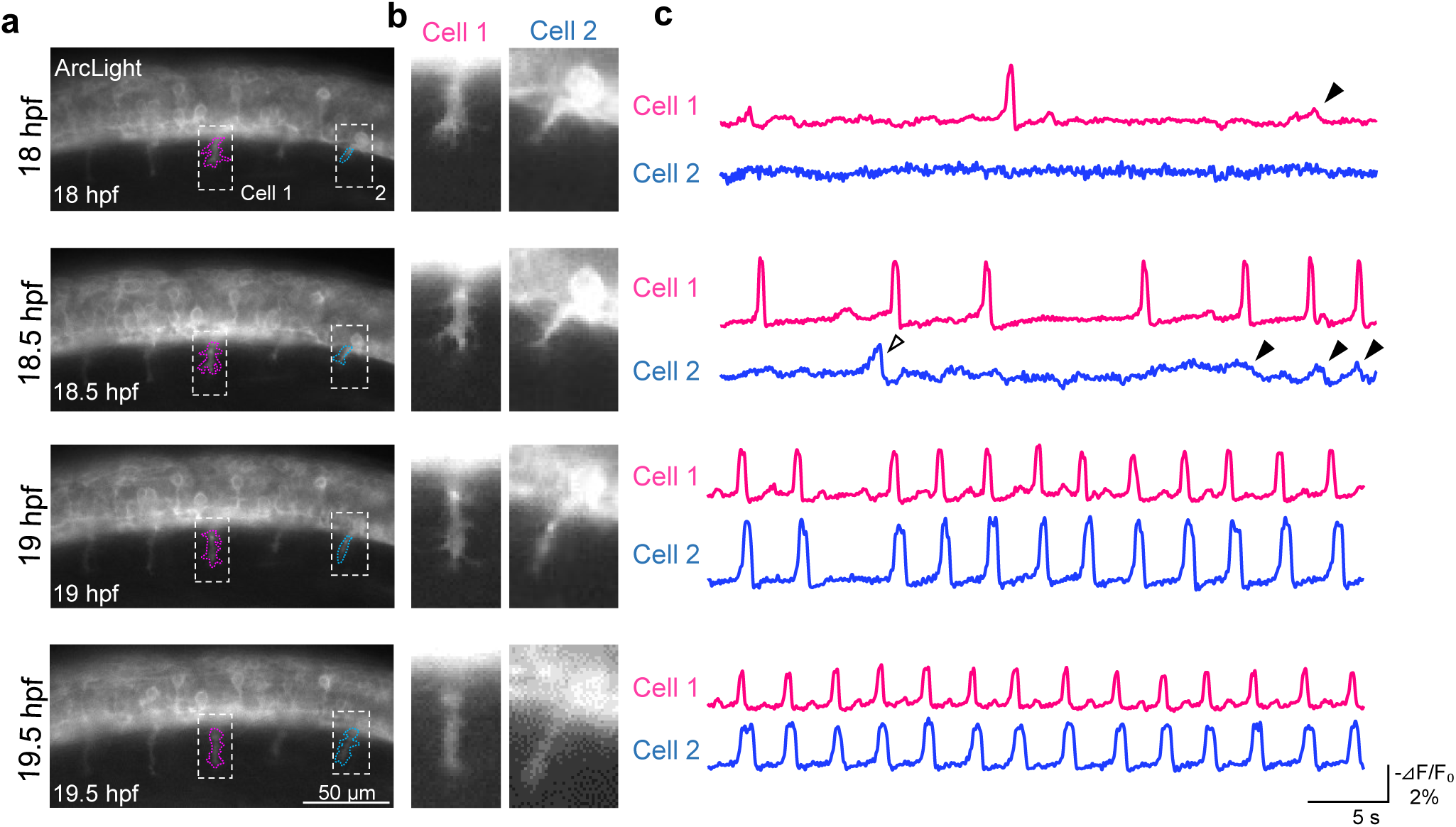
Long-term voltage imaging of the spinal cord PMNs shows the emergence of population activity. (a) Left views of the spinal cord of ArcLight positive fish from 18 hpf to 19.5 hpf. (b) Enlarged views of (a). (c) Voltage dynamics obtained from the axons of the two cells shown in (a). Black and white arrowheads indicate small depolarization and immature firing, respectively.

**Figure 6.**
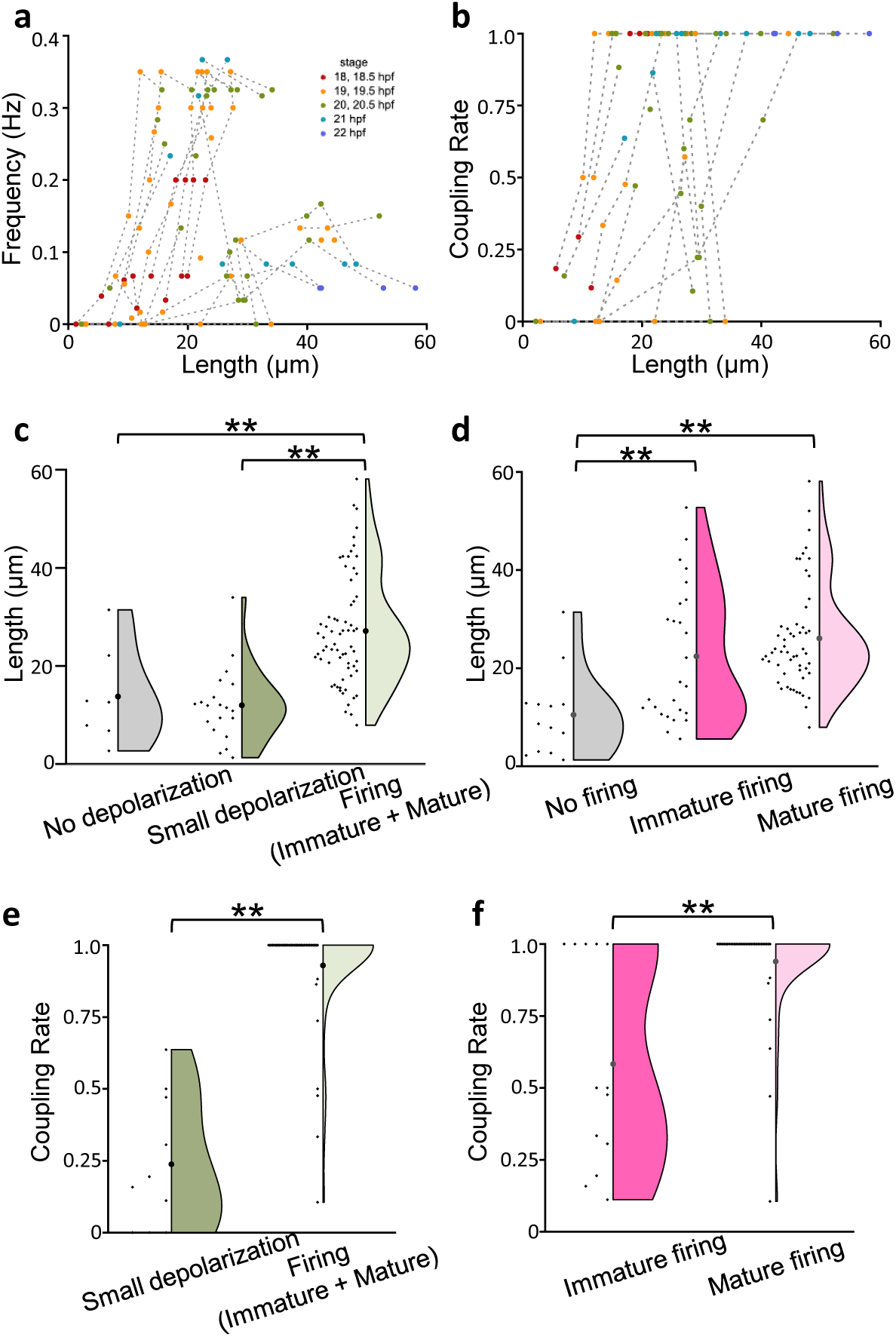
Characteristic patterns of depolarization observed during the development of PMNs population activity. Quantitative analysis indicates that coupled activity among PMNs emerged via small depolarization and immature firing. (a,b) (a) The relationship between axon length and firing frequency. (b) The relationship between axon length and coordination with neighboring neurons. Developmental process of each PMN is indicated by lines. Developmental stages are indicated by dots colors. (c) The relationship between the axonal length and existence of small depolarization (no depolarization: 13.76±3.75 μm, small depolarization: 11.97±1.67 μm, Firing: 27.12±1.38 μm, 5 fish, 30 cells, **p<0.01, Mann–Whitney U test). (d) The relationship between axonal length and immature firings (no depolarization: 10.48±2.38 μm, small depolarization: 22.42±2.69 μm, Firing: 26.06±1.44 μm, 5 fish, 30 cells, **p<0.01, Mann–Whitney U test). (e) Comparison of coupling rates between PMNs with small depolarizations and those with firings (immature and mature firings) (small depolarization: 0.24±0.07, firing: 0.93±0.03, 4 fish, 18 cells,, **p<0.01, Mann– Whitney U test). (f) Comparison of coupling rates between PMNs exhibiting immature and mature firings (immature firing: 0.58±0.10, mature firing: 0.94±0.03, 4 fish, 18 cells, **p<0.01, Mann–Whitney U test).

To further assess how spontaneous activity evolves later in development, we extended voltage imaging to 23 hpf. By this stage, the previously regular and rapid firing observed at 20 hpf had decreased (Fig. 7, Supplementary Fig. S8). Depolarization patterns also changed—bursts became longer and left–right alternation diminished (Fig. 7b). Cross-correlation analysis revealed that at 20 hpf, neurons on both sides of the spinal cord were highly synchronized and exhibited strong periodicity (Fig. 7c). By 23 hpf, this coordination was largely diminished, although synchrony among ipsilateral neurons persisted. Notably, left–right alternating activity, which was prominent at 20–21 hpf, was markedly reduced at 23–24 hpf (Fig. 7d). These results suggest that while coordinated PMN activity declines during later stages of development, intra-hemispheric synchrony is maintained.

**Figure 7.**
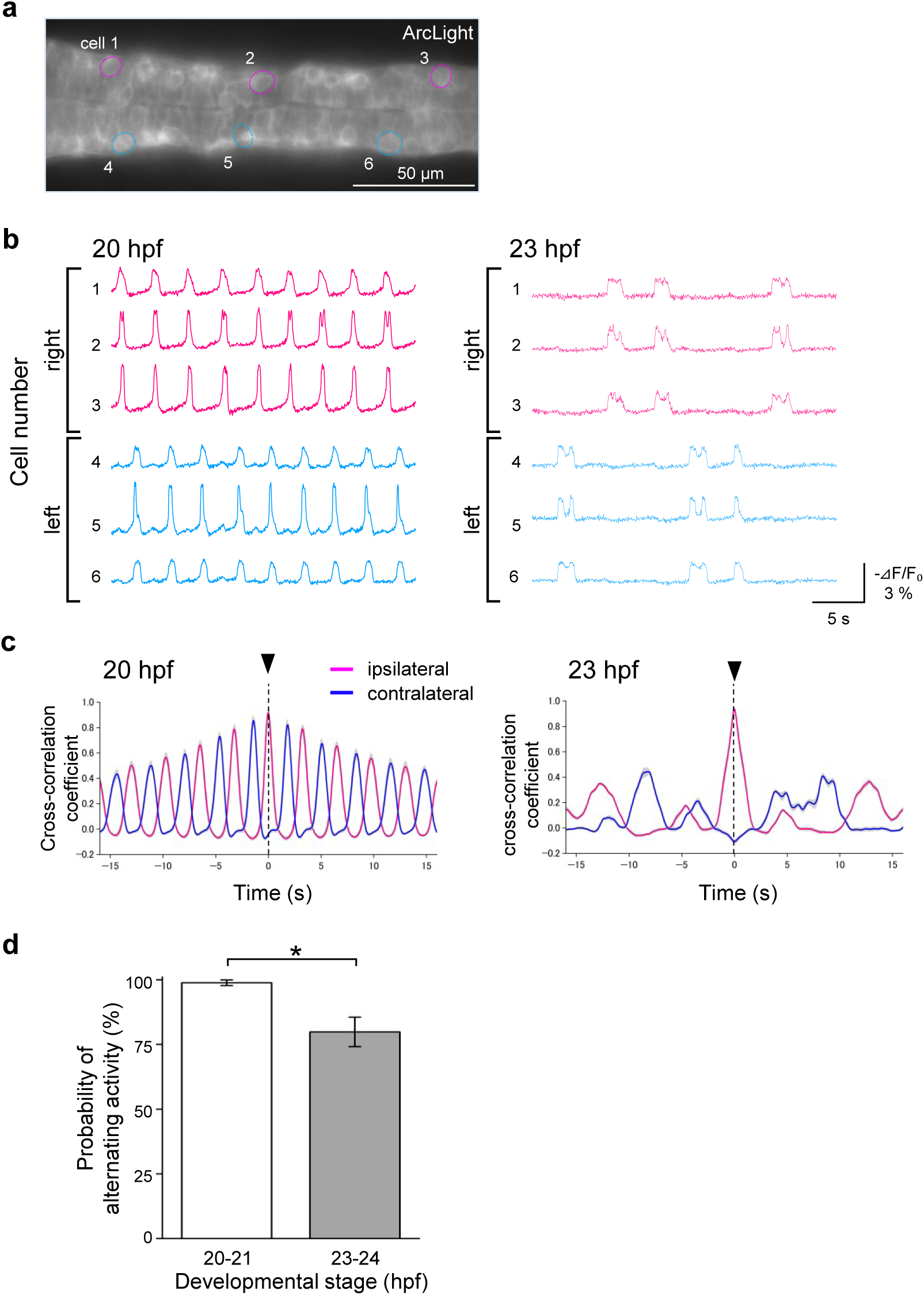
Changes in firing properties in spinal cord neuron populations. (a) Dorsal view of ArcLight positive spinal cord at 20 hpf. (b) Voltage dynamics of each neuron are shown at 20 and 23 hpf. The position of ROIs is indicated in (a). (c) Cross-correlation analysis of ipsilateral and contralateral pairs of spinal cord neurons at 20 and 23 hpf. (d) Probability of altering activity at 23-24 hpf was significantly reduced compared to that at 23-24 hpf. 20–21 hpf: 98.9±1.10%; 23–24 hpf: 98.9±1.10%, 5 fish; 23–24 hpf: 79.8%±5.71%, 4 fish; p<0.05, Mann–Whitney U test.

## Discussion

Understanding how neuronal populations generate coordinated activity during development is a central challenge in neuroscience. Electrophysiological recordings have historically provided valuable insights into the membrane dynamics of spinal neurons. However, these techniques are invasive and largely limited to short-duration and single-cell level [43, 46]. Calcium imaging has been used to examine developing circuits but lacks the temporal resolution and direct voltage sensitivity required to resolve fine membrane potential dynamics [42, 47, 48]. In this context, genetically encoded voltage indicators (GEVIs) offer a promising alternative, providing direct, fast, and non-invasive readouts of membrane potential. While GEVIs have been applied in mature brain tissues and cardiac tissues, their application to vertebrate embryonic development has remained rare due to technical limitations. To the best of our knowledge, this is the first study to achieve long-term voltage imaging of genetically defined neuronal populations in the spinal cord during development in a vertebrate model.

### High-resolution imaging of voltage and morphological dynamics

Subcellular compartments—such as dendrites axons, and filopodia—play essential roles in neuronal function, shaping synaptic integration and signal propagation [44, 45]. In this study, ArcLight enabled voltage imaging not only at the soma, but also at proximal and distal axonal regions of primary motor neurons (PMNs). Notably, voltage fluctuations were also observed in filopodia, suggesting that even fine structures can exhibit membrane potential dynamics. Such fine-scale resolution opens opportunities to investigate the bioelectrical properties of neurite outgrowth, synaptogenesis, and structural remodeling. Combining voltage imaging with ratiometric or dual-color imaging approaches may allow simultaneous measurement of voltage and morphological parameters at high spatiotemporal precision [49].

### Long-term population-level voltage dynamics in the developing spinal cord

The zebrafish spinal cord possesses one of the earliest neuronal circuits to form during development, and its transparency makes it an ideal model for in vivo imaging. Using a newly generated ArcLight transgenic line, we visualized voltage dynamics in spinal neuron populations from early developmental stages. At 20 hpf, we detected left-right alternating depolarizations, consistent with periodic depolarizations (PDs) previously observed via electrophysiology [34]. Compared to calcium imaging or earlier voltage sensors like ASAP1, ArcLight offered superior temporal resolution and signal-to-noise ratio, respectively [13, 50]. This enabled detection of not only suprathreshold PDs, but also subthreshold events and hyperpolarizations, capturing a wider dynamic range of neuronal activity. Importantly, ArcLight could resolve activity patterns at both the single-cell and population levels, providing a comprehensive picture of how spinal circuits operate and evolve during development.

### Emergence of voltage dynamics in the spinal neuron populations

One of the significant findings of this study is the stepwise emergence of coordinated voltage dynamics in motor neurons, closely tied to their morphological maturation. By performing long-term voltage imaging from, we found that PMNs followed a developmental sequence: (1) absence of depolarization, (2) emergence of small voltage fluctuations, (3) irregular and weakly coordinated activity, and (4) synchronized, robust left-right firing. This progression was strongly correlated with axonal extension, suggesting a tight coupling between structure and function.

It has been implicated that spinal cord neurons do not fire before axonal outgrowth, but detailed observation was missing [46]. Interestingly, here early voltage fluctuations preceded axon outgrowth, possibly driven by intrinsic dynamics of ion channels or electrical coupling via gap junctions, which predominate at these early stages. These small fluctuations may represent primitive forms of population synchrony, preceding the establishment of mature synaptic connections. This idea is consistent with other model systems where early electrical coupling plays an instructive role in circuit patterning [51].

Our approach provides a first-in-class demonstration of long-term, non-invasive voltage imaging in a developing vertebrate nervous system. Beyond the spinal cord, this approach can be extended to other regions, enabling a detailed dissection of diverse population dynamics during development. Furthermore, applying this imaging system to genetic models of neurodevelopmental disorders— such as autism spectrum disorders (ASD), epilepsy, or motor neuron disease—may allow the identification of early disruptions in coordinated neuronal activity. Detecting such early electrophysiological phenotypes could contribute to a better understanding of disease mechanisms.

In conclusion, we have established GEVI-based long-term voltage imaging platform for long-term, high-resolution monitoring of developing neuronal populations in vivo. This system revealed the temporal emergence and maturation of population-level voltage dynamics in the spinal cord, offering new insights into how coordinated neural activity is built during development. The combination of spatial, temporal, and functional resolution achieved here represents a critical advance for the field of developmental neuroscience. By integrating this approach with optogenetics, cell-type-specific reporters, or functional perturbation, future studies can address how intrinsic and extrinsic factors shape circuit assembly. Moreover, our findings highlight the value of voltage imaging as a general tool for probing bioelectric mechanisms in morphogenesis, circuit development, and disease [52, 53].

## Acknowledgement

We thank Drs Koichi Kawakami, Shinichi Higashijima, Hitoshi Okamoto, Hisashi Kakinuma, and Ryunosuke Amo for providing fish lines; Kyo Yamasu for plasmid DNA; Junichi Nakai, Masamichi Ohkura, and Keiko Ando for technical support; and Lawrence Cohen and Shinichi Higashijima for helpful discussions. This work was supported in part by JSPS KAKENHI (Grants-in-Aid for Scientific Research), JST Tenure Track program at Saitama University (SUTT), Strategic Research Program for Brain Science from Ministry of Education, Culture, Sports, JST A-STEP Grant Number JPMJTM20C05, New Energy and Industrial Technology Development Organization (NEDO), Kao Foundation grant, Asahi Glass Foundation, and Kato-bio Foundation.

## Author Contributions

A.S, A.H., N.F., M.H., K.M., and S.T. performed the experiments. S.T. conceived and designed the experiments. S.T., A.S., A.H., and N.F. wrote the manuscript. All authors reviewed the manuscript.

---

**Supplementary Figure S1**

Monitoring spontaneous activity of individual cells in the spinal cord by calcium imaging by GCaMP6f. (a) Fluorescence image of the ventral spinal cord of Tg*(elavl3: GCaMP6f)* embryo at 21 hpf. ROIs were placed on six cells (red: right side, blue: left side). Higher-magnification views are shown on the right. (b) Schematic diagram of the recording setup. (c) Activity patterns of the six cells shown in (a). (d) A higher magnification view of the ArcLight signal of cells 1 and 5. (e) Simultaneous recordings of membrane potential by cell-attached patch-clamp recordings and ArcLight imaging showed that the recorded neurons were indeed depolarized. A fluorescence image of the spinal cord of Tg*(HuC:GVP;UAS:ArcLight)* fish at 23 hpf. (f) Voltage (top) and optical (bottom) signals detected at the cell indicated in (e).

**Supplementary Figure S2**

Confocal observation of ArcLight expression and voltage dynamics in the spinal cord. (a) Lateral views of Tg*(elavl3:GAL4-VP16;UAS:ArcLight)* embryo at 1 dpf. Higher magnification images shown on the right indicate filopodia and lamellipodia of primary motor neurons (arrowheads). (b) Voltage imaging of cell1 in (a). ROIs were set at the axonal region. ArcLight detected depolarization at axons (black arrowheads). The gray area is the period when the embryos moved.

**Supplementary Figure S3**

Voltage imaging of a primary motor neuron at higher spatial resolution. (a) Left views of the spinal cord at two different time points. Arrowheads indicate filopodia (white, yellow). A yellow arrowhead indicates a filopodium that shows a prominent extension. (b, c) An example of voltage dynamics detected at various positions in the axon. ROI4-6 were set on a filopodium. ROI 7 was set in a region where no clear structure was observed.

**Supplementary Figure S4**

Single-cell labeling by ArcLight shows the morphological changes of the spinal cord neurons. Images taken at 21 hpf and 22 hpf are shown. An arrowhead indicates a growth cone of the extending axon.

**Supplementary Figure S5**

Developmental changes in the voltage dynamics of PMNs at the soma and axons. Voltage dynamics were generally similar between the soma and axons. Representative traces from Cell 2 in Figure 5 are shown.

**Supplementary Figure S6**

Comparison of voltage dynamics between immature and mature firings. Immature firings showed a significantly longer time to peak (a) but not decay time (b) (time to peak; immature firing: 1.31±0.43 s, mature firing: 0.63±0.20 s, decay time; immature firing: 0.59±0.25 s, mature firing: 0.50±0.17 s, **p < 0.01, Mann–Whitney U test, 3 fish, 65 transients).

**Supplementary Figure S7**

Developmental change in the voltage dynamics of PMNs. (a-e) Left-side views of the spinal cord of ArcLight-positive fish from 18 to 20 hpf (top), and corresponding voltage traces (bottom) of the PMNs indicated by arrowheads. ROIs were set on axons, as shown in (f).

**Supplementary Figure S8**

Long-term voltage imaging shows the changes in the firing patterns of the spinal cord neurons population. (a) ROIs position of cells 1-6. (b) Voltage dynamics of the 6 cells are shown from 20 to 23 hpf. Cells on the right and left sides of the spinal cord are shown in red and blue, respectively.

**Supplementary Movie1**

Spontaneous activity of individual cells in the spinal cord is observed by ArcLight. Dorsal view of the ventral spinal cord of Tg*(elavl3:GAL4-VP16;UAS:ArcLight)* fish at 21 hpf.

**Supplementary Movie2**

Monitoring of spontaneous activity of spinal cord neurons, including primary motor neurons by ArcLight. Left view of the spinal cord of Tg*(elavl3:GAL4-VP16;UAS:ArcLight)* fish at 20 hpf.

**Supplementary Movie3**

Single-cell labeling and voltage imaging of spinal cord neurons using ArcLight. Left view of the spinal cord of Tg*(elavl3:GAL4-VP16)* embryo at 21 hpf. A high-resolution image is shown in Supplementary Figure S4.

**Supplementary Movie4**

Voltage dynamics of an axonal region of a primary motor neuron at a high spatial resolution. Left view of the spinal cord Tg*(elavl3:GAL4-VP16;UAS:ArcLight)* fish at 18.5 hpf.

**Supplementary Movie5**

Voltage imaging of spinal neurons at 18.5 hpf (12 s). Left-side view is shown.

**Supplementary Movie6**

Voltage imaging of spinal neurons at 19 hpf (12 s). Left-side view is shown.

**Supplementary Movie7**

Voltage imaging of spinal neurons at 19.5 hpf (12 s). Left-side view is shown.

